# Enhanced protein secretion in reduced genome strains of *Streptomyces lividans*

**DOI:** 10.1101/2023.02.14.528591

**Authors:** M. B. Hamed, T. Busche, K. Simoens, S. Carpentier, J. Kormanec, L. Van Mellaert, J. Anné, J. Kalinowski, K. Bernaerts, S. Karamanou, A. Economou

## Abstract

*S. lividans* TK24 is a popular host for the production of small molecules and for the secretion of heterologous proteins. TK24 has a large genome with at least 29 secondary metabolite gene clusters that are non-essential for viability and undergo complex regulation. To optimize heterologous protein secretion, we previously constructed ten chassis strains that are devoid of several secondary metabolite gene clusters. Genome reduction was aimed at reducing carbon flow to secondary metabolites and pigmentation in the spent growth medium and improving colony morphology. Strains RG1.0-RG1.10 contain various deletion combinations of the blue actinorhodin cluster (*act*), the calcium-dependent antibiotic (*cda*), the undecylprodigiosin (*red*) and coelimycin A (*cpk*) clusters, the melanin cluster (*mel)*, the *mat*AB genes that affect mycelial aggregation and the non-essential sigma factor *hrd*D that controls the transcription of Act and Red regulatory proteins. Two derivative strains, RG1.5 and 1.9, showed a ∼15% reduction in growth rate, >2-fold increase in the total mass yield of their native secretome and altered abundance of several specific proteins compared with TK24. Metabolomics and RNAseq analysis revealed that genome reduction led to rapid cessation of growth due to aminoacid depletion and caused both redox and cell envelope stresses, upregulation of the Sec-pathway components *secDF* and chaperones and a cell envelope two component regulator. RG1.9 maintained elevated heterologous secretion of mRFP and mTNFα by 12-70%. An integrated model is presented linking genome reduction and enhanced secretion.

## Introduction

*Streptomyces* is a ubiquitous soil organism, known as a copious secretor of polypeptides and secondary metabolites (1, 2). The lack of an outer membrane and a relatively low level of secreted proteases makes it an attractive platform for the production/secretion of heterologous polypeptides of bacterial and eukaryotic origin (3). *Streptomyces lividans* TK24 has been successfully used as a host for the production of several heterologous proteins such as: active trimeric murine tumor necrosis factor-alpha (mTNF-α)(4, 5), human tumor necrosis factor (6), *Jonesia sp.* xyloglucanase (7), a thermostable cellulase (CelA) (2), a *Thermobifida fusca* endoglucancase (8), a *Streptoverticillium cinnamoneum* phospholipase D (9), an *Actinoallomurus sp.* glutenase (10), an *Aeropyrum pernix* pernisine (11, 12), a *S. halstedii* phospholipase D (13) and a monomeric red fluorescent protein (mRFP) (3).

The prodigious production of secondary metabolites such as actinorhodin (*act*) and undecylprodigiosin (*red*) is controlled by environmental and physiological conditions, such as starvation from nutrients like phosphate (14). Secondary metabolites are synthesized by the peptide (15), the polyketide synthase (PKS) (16), the non-ribosomal polypeptide synthase (NRPS) (17), the shikimate (18), the β-lactam synthesis (19), and the carbohydrate (15) pathways. The genes for these biosynthetic pathways are mostly clustered in the genome and regulated by pathway-specific transcriptional activators termed “cluster-situated regulators (CSR)” encoded by genes within these clusters (20, 21). The number of secondary metabolic gene clusters varies between *Streptomyces* species, e.g. 37 in *S. avermitilis* (22), 29 in *S. lividans* TK24 and *S. coelicolor* A3 (2) (22–24). The chemical structure of <30% of the synthesized compounds has been elucidated (25). They belong to various groups: polyketides, pyrones, peptides, siderophores, γ-butyrolactones, butenolides, furans, terpenoids, fatty acids, oligopyrroles, and deoxysugars. Most studies have focused on secondary metabolite clusters in *Streptomyces* that encode the polyketide-derived antibiotics *act*, coelimycin A (*cpk* A, precursor of yellow coelimycins P1 and P2)), *red*, and a calcium-dependent ionophore antibiotic (*cda*) as they are the endogenous highly produced antibiotics and they have roles in cell differentiation (26, 27) (22).

*Streptomyces* antibiotic regulatory proteins (SARPs) are the most frequently encountered CSRs among the *Streptomycetes* (28). SARPs for the most studied clusters are ActII-ORF4, RedD, CdaR, and CpKO for *act*, *red*, *cda* and *cpk*, respectively (22). The transcription of ActII-ORF4 and RedD in *S. coelicolor* A3(2) is initiated by an RNA polymerase holoenzyme containing the non-essential *hrdD* sigma factor (SCO3202) (29–31). The expression of SARPs was upregulated in conjunction with morphological or physiological differentiation (32). Thus in *S. coelicolor* transcription of *redD* and *cdaR* increased more than two-fold before the late-exponential phase, while the *act*II-*ORF4* gene showed elevated transcription before stationary phase (33).

Reduced genome strains, that contain essential genes but are missing increasing numbers of genes considered to be “redundant” have been designed in many bacteria and resulted in some cases in unexpected phenotypes such as growth rate reduction (34). These genome reduction efforts mainly aimed at improving the performance of bacteria by enhancing their metabolic efficiency and decreasing the redundancy in genes and regulatory circuits (35, 36). For example, JCVI-syn3.0, a designed *Mycoplasma mycoides* strain, contains only essential genes and showed impaired growth (37). In *E. coli* a series of strains with genomes deleted of non-essential genes was constructed in a step-wise manner (from 48-982 kb). The derivative strains showed a positive correlation between genome shortening and reduction in growth rate (38). On the other hand, 12 K-islands, which contain transposon, cryptic phage, damaged, and genes of unknown-function, representing 8.1% of the *E. coli* K-12 MG1655 genome were deleted without any effect on growth (39). Moreover, 1 Mbp was deleted from *E. coli* K-12 to produce MGF-01 that showed better growth compared with the parent strain (40).

In *Streptomyces*, there have been many attempts to construct an optimized strain with an efficient unperturbed supply of precursors for the optimal biosynthesis of target heterologous proteins and antibiotics (26, 41, 42). This was generally executed by deleting specific protease and secondary metabolic genes (26, 30, 42, 43). For example deletion of the gene for the ATP-dependent zinc metalloprotease FtsH in *S. lividans* TK24 reduced the dry cell mass by 37% while it elevated the expression and secretion of heterologous mRFP by 29-fold compared with the wild type (42). Similarly, reintroduced actinorhodin production was enhanced in *S. coelicolor* missing 10 polyketide and non-ribosomal peptide gene clusters (44); both streptomycin and cephamycin C were efficiently produced in *S. avermitilis* with >1.4 Mb deletion (45); heterologous production of chloramphenicol and congocidine were significantly increased in *S. coelicolor* M145 that had lost the *act*, *red*, *cpk* and *cda* clusters (27); the antitumoral polyketide mithramycin A was produced at 2-fold higher yields in a *S. lividans* TK24 derivative that lacked the *cpk*, *act*, *red* and *cda* clusters than in the wild type (43).

Bacterial genome mining, to create a simplified cell with predictable behaviour, positively affected the protein production yield. In Gram-positive *Bacillus subtilis*, deletion of 11 non-essential gene clusters (865 genes) increased the secretion of heterologous cellulase and M-protease by 70 and 150%, respectively (46). Yet another, reduction of the *B. subtilis* genome by 20.7% increased the production of secreted alkaline cellulase Egl237 by ∼2-fold (47). Deletion of 2.8% of the *Lactococcus lactis* NZ9000 genome (including prophages, transposons, and related proteins), elevated the expression and secretion of red fluorescent protein by >2 fold (48).

To construct a versatile, simple reduced genome chassis of *S. lividans* TK24, five secondary metabolite genes clusters and three individual genes were previously deleted in different combinations to generate 10 strains (RG1.0 to RG1.10) (Fig. 1; Table 1 and S1) (41, 43). The deleted clusters encoded for four pigmented antibiotics: *act* (26.7 kb), *red* (39.3 kb), *cpk* (59.1 kb) *mel* (8.9 kb), and *cda* (26.7 kb). The extracellular glycan encoding genes *matA* and *matB*, and the region encoding transcription factor *hrdD* were also selected (Fig. 1). These regions/genes were selected based on one of the following criteria: (I) their biosynthesis is associated with cell differentiation or autolysis; (II) the ability to interact with heterologous secreted proteins that affect their industrial bioprocessing, e.g. *mel* (43, 49); (III) a role in cell aggregation; (VI) they showed high transcription levels in minimal medium.

**Fig. 1.**
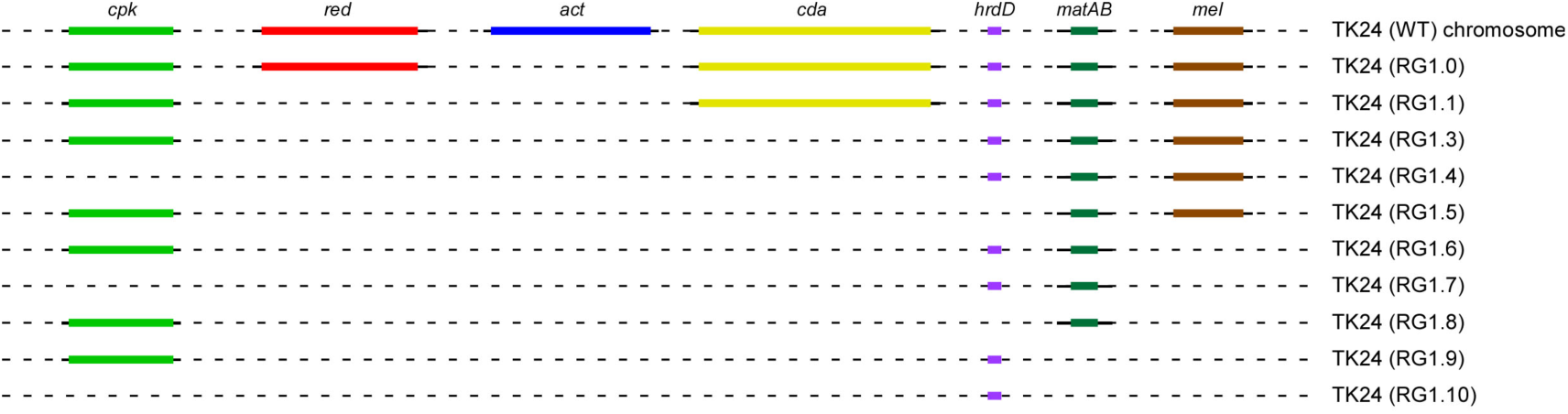
Deleted secondary metabolite gene cluster in *S. lividans* TK24. Physical maps of the chromosome regions of *S. lividans* TK24 corresponding to the actinorhodin biosynthetic gene cluster (26.7 kb), undecylprodigiosine biosynthetic gene cluster (39.3 kb), the calcium dependent antibiotic gene cluster (26.7 kb), region encoding transcription factor HrdD (1.3 kb), coelimycin P1 gene cluster (59.1 kb), melanin gene cluster (8.9 kb) and aggregation genes *mat*A and *mat*B (4.5 kb), which were deleted.

**Table 1:**
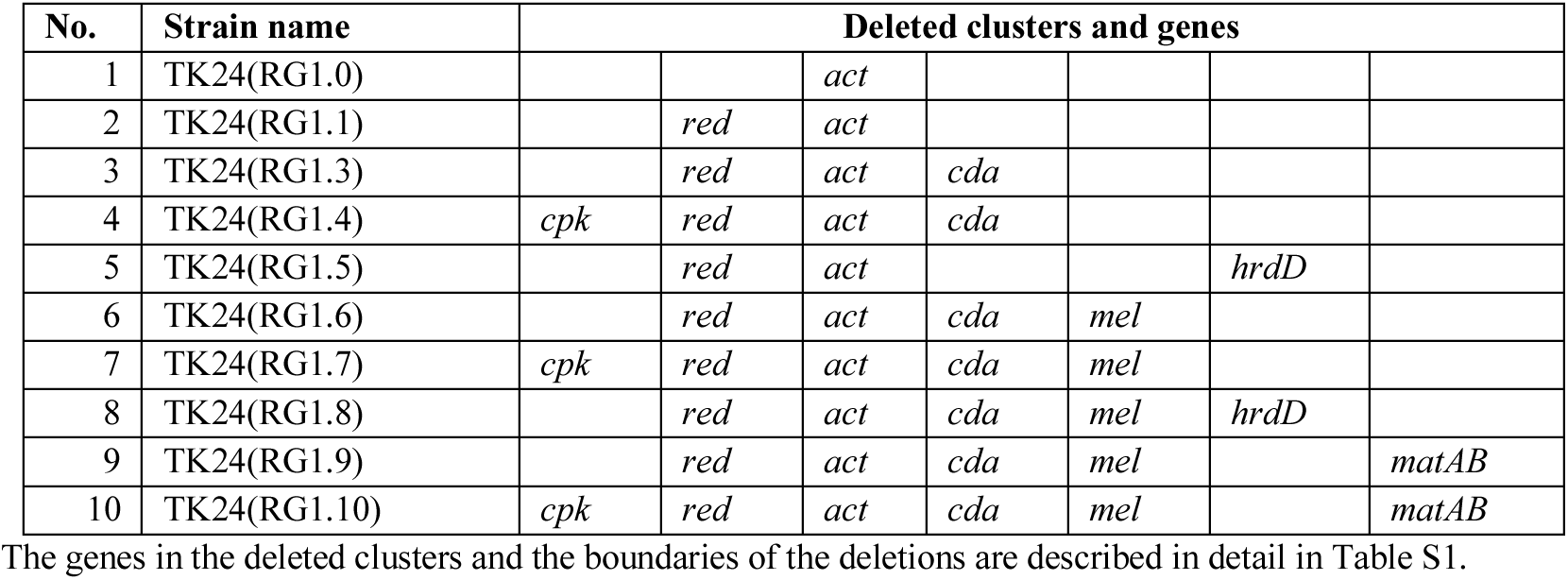
Deletion of secondary metabolite gene clusters in the reduced genome strains

| No. | Strain name | Deleted clusters and genes |  |  |  |  |  |  |
| --- | --- | --- | --- | --- | --- | --- | --- | --- |
| 1 | TK24(RG1.0) |  |  | *act* |  |  |  |  |
| 2 | TK24(RG1.1) |  | *red* | *act* |  |  |  |  |
| 3 | TK24(RG1.3) |  | *red* | *act* | *cda* |  |  |  |
| 4 | TK24(RG1.4) | *cpk* | *red* | *act* | *cda* |  |  |  |
| 5 | TK24(RG1.5) |  | *red* | *act* |  |  | *hrdD* |  |
| 6 | TK24(RG1.6) |  | *red* | *act* | *cda* | *mel* |  |  |
| 7 | TK24(RG1.7) | *cpk* | *red* | *act* | *cda* | *mel* |  |  |
| 8 | TK24(RG1.8) |  | *red* | *act* | *cda* | *mel* | *hrdD* |  |
| 9 | TK24(RG1.9) |  | *red* | *act* | *cda* | *mel* |  | *matAB* |
| 10 | TK24(RG1.10) | *cpk* | *red* | *act* | *cda* | *mel* |  | *matAB* |

The genes in the deleted clusters and the boundaries of the deletions are described in detail in Table S1.

The mutated strains showed differences in their total native secreted proteome compared with TK24. RG1.9 produced native secretome amounts >2 times more than the wild type that contained 86 differentially abundant proteins compared with the parent strain, respectively, with more secreted proteins in RG1.9. Transcriptomics analysis of RG1.9 showed high expression of the gene encoding for *secDF*, four cytoplasmic chaperones, a two-component system (*SLIV_09695* and *SLIV_09690*) and two sigma factors *sig^Q^* (SLIV_13835) and *sig^70^* (SLIV_13900). The upregulation of this two-component system is positively-correlated with overproduction of extracellular proteins (50). Additionally, RG1.9 showed upregulation for genes related to both redox and cell envelope stresses and four genes related to the arginine biosynthesis pathway. Moreover, aspartate, glutamate and asparagine were depleted after the mid-exponential phase that caused suppression in the growth rate. The elevated native secretome levels reflected an overall improved capacity of these strains for generic protein secretion seen with the secretion of the heterologous proteins mRFP (3), and mTNF (5) that were higher by 12-70% in RG1.9 than in the wild-type TK24 strain.

This study paves the way to understanding how secondary metabolite clusters can be manipulated to construct a surrogate TK24 platform with optimized metabolite funnelling to heterologous secretion.

## Results

### Transcription of secondary metabolite gene clusters in *S. lividans* TK24

Prior to any downstream analysis, the transcription levels of the selected genes and gene clusters deleted in the reduced genome strains (RG1.0 to RG1.10) (Fig. 1 and S1; Table 1) were analyzed. *S. lividans* TK24 was grown in a minimal medium supplemented with glucose, a traditional medium for streptomycetes growth. Cells were collected at three different growth phases (early, late-exponential and stationary) and the transcriptomes of these cells were analysed using RNAseq. Under these conditions, the transcript levels for most of the secondary metabolite gene clusters, including those deleted during genome reduction, were high at the late-logarithmic phase (Fig. S1). The *act* cluster (*SLIV_12925* to *SLIV_13030*) showed the highest transcript levels in both late-exponential and stationary phases (Fig. S1). Similarly, secondary metabolite regulatory proteins (SARPs), antibiotics cluster activators (51), *red*Z (*SLIV_09200*), *red*D (*SLIV_09220*), *cda*R (*SLIV_21605*) showed high transcript levels in the late-log phase (Fig. S1; bold red; Table S2). Moreover, *act*II-*ORF4* (*SLIV_12960*) showed the highest transcript levels in both late-exponential and stationary phases (Fig. S1; bold red; Table S2). On the other hand, *cpk*O (*SLIV_06745*) and *cpk*N (*SLIV_06705*), showed moderate transcript levels during all the growth phases (Fig. S1; bold and red; Table S2). The upregulation of SARPs is consistent with what was previously reported in *S. coelicolor* A3(2) (52).

### Effect of deleted clusters on cell growth and native secretome production

The effect of cluster and gene deletions on cell growth was tested for all the engineered strains. The strains were grown in nutrient broth (NB) medium that has been previously described as the best-performing medium for native and heterologous protein production (2, 3, 42). The strains were inoculated in NB based on their growth at OD_600_ in order to have equal amounts of mycelia for each strain (see Materials and methods). Most strains showed similar growth patterns compared with TK24 for both the late-exponential and stationary phase by comparing the dry cell masses (g/L) of the strains (Fig. S2A). RG1.1 and RG1.7 produce 40% more and RG1.5 and RG1.9 15% less dry cell weight than the wild type, respectively (Fig. 2A and S2A).

**Fig. 2.**
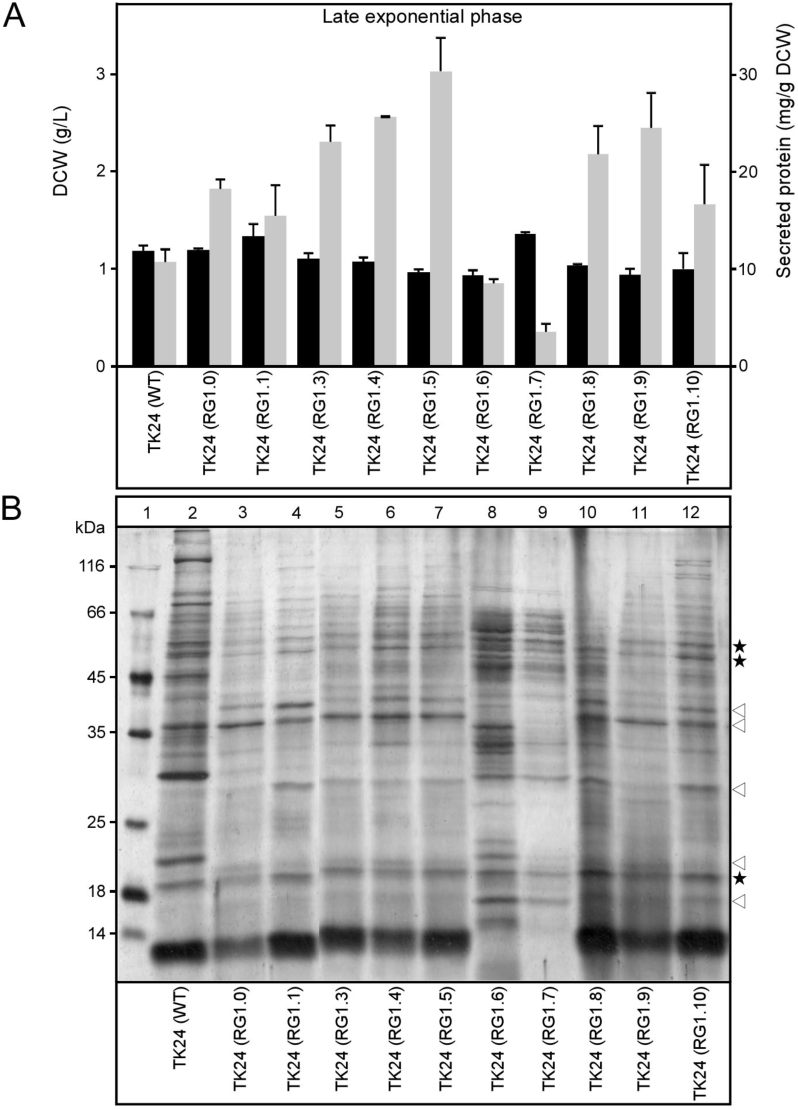
Reduced genome strains cell growth and native secretome production. **(A)** Comparison of total native secretome produced in (mg) (shown as grey bars) correlated to a gram of dry cell weight (DCW) and the dry cell weight produced in (g/L) (shown as black bars) for Reduced genome strains compared with *S. lividans* TK24 in nutrient broth medium (NB) at the late exponential phase. *n*=3, values represent the mean ± SD. **(B)** Polypeptides (3 µg/lane) from culture supernatants (2-16 µL/lane) of TK24 or the reduced genome strains grown for the indicated times in nutrient broth (NB) were analyzed by SDS-PAGE and silver staining. Lane 1, molecular weight markers: β-galactosidase (116 kDa), bovine serum albumin (66.2 kDa), ovalbumin (45 kDa), lactate dehydrogenase (35 kDa), restriction endonuclease Bsp98I (25 kDa), β-lactoglobulin (18.4 kDa), lysozyme (14.4 kDa). Asterisks: secretome polypeptides with minor changes; empty arrows: secretome polypeptides with major changes.

To determine the effect of genome deletions on the production of the native secretome, the amounts of total native secretome produced by these strains was compared to those of a gram of dry cell weight (DCW) of TK24 (Fig. 2A). Most strains produced more secretome mass than TK24. The highest amount of the total secretome was obtained with RG1.5, which was three times more (∼ 30 mg/g DCW). Other strains (RG1.3, 1.4, 1.8 and 1.9) produced twice more. On the other hand, two strains (RG1.6 and RG1.7) secreted less total polypeptide mass than the WT (Fig. 2A). A negative correlation of the slow growth rate of cells and total protein secreted observed in some of the mutated strains matches what we previously reported (2, 3, 53). Apparently, excessive secretion is stimulated when carbon backbones cannot be metabolically funneled towards cell growth.

To study the effect of the deleted clusters on the native secretome profile, the total secreted polypeptides of TK24 and its reduced genome strains were grown in NB medium for 24h, and the whole supernatant was harvested and analyzed by SDS-PAGE and silver staining (Fig. 2B). The protein patterns obtained varied between the mutated strains and those of the wild type. Several different new polypeptides appeared while others were lost (Fig. 2B). The abundance of some proteins was similar for the mutated strains and TK24 (Fig. 2B; asterisks), while that of other proteins changed in the secretomes of some RG strains (Fig. 2B; empty arrows).

To determine the altered secretomes in depth, we analyzed them by label-free nanoLC-MS/MS. We examined the secretome of three mutated strains (RG1.4, RG1.5 and RG1.9) which produced the highest amount of native secretome in comparison with the parent *S. lividans* TK24 (Fig. 2A). The secreted polypeptides were expressed per unit of dry cell mass of the mutated strains or the wild type. These polypeptides were then identified and quantified by MS (Fig. 3 and S3; Table S3 and S4). The abundance of proteins in the secretome of the mutated strains was compared to that of the wild type (Fig. 3 and S3). In all cases, several polypeptides, representing ∼10-15% of the secretome, were identified at abundances different to those of the wild type strain. The abundances of 62, 55 and 86 secreted proteins were statistically different in RG1.4, RG1.5 and RG1.9 compared with the wild type, respectively (Fig. 3A and S3A). Out of these, 39 and 36 proteins were more abundant in the RG1.4 and RG1.5 secretomes compared with TK24, respectively, while 67 proteins were more abundant in the RG1.9 secretome, including proteins that are seen at both lower and higher levels in the mutant strain (Fig. 3 and S3; Table S4). Secreted proteins whose levels were affected include: a putative lipoprotein localization LolA/LolB/LppX-like protein and N-acetylmuramoyl-L-alanine amidase that were synthesized/secreted in all of the mutated strains >1.6-3 times more than in TK24. In contrast, phospholipase A2 (D6ES51) and A0A076MBH5 of unknown function secreted through the Tat pathway, were secreted > 1.5-6-fold less.

**Fig. 3:**
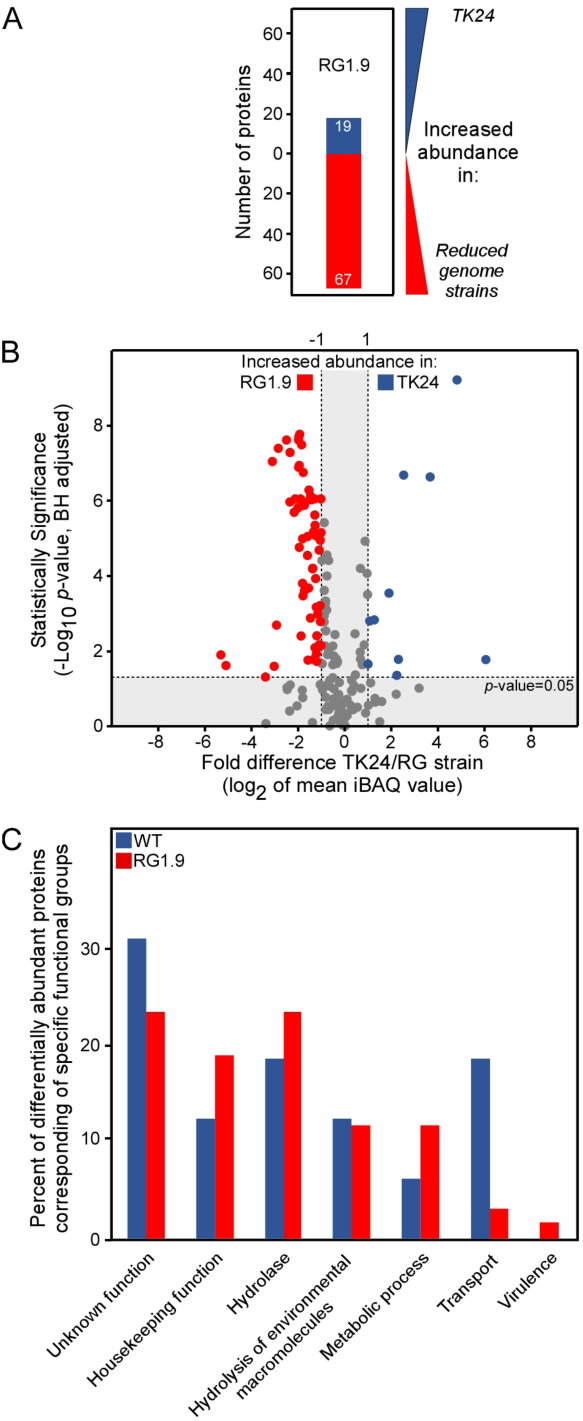
Comparative secretome analysis of the reduced genome strains against TK24. **(A)** Number of differentially abundant secreted proteins between RG1.9 against the WT. Proteins with increased abundance in WT are shown in blue and proteins more abundant in the mutated strain are colored red. Samples were loaded for proteomic analysis normalized to the same amount of cell biomass. **(B)** Volcano-plots showing the differentially abundant proteins between TK24 and reduced genome strain RG1.9. Each dot represents one protein. On the *x* axis is plotted the fold difference (in log2 scale) of the mean protein abundance in the TK24 (marked: WT) over that in RG1.9, and on the *y* axis the *p*-value derived from a *t*-test between the two strains (–log10, adjusted by the Benjamini–Hochberg method). With blue are colored proteins more abundant in the WT and in red, the significantly more abundant proteins in RG1.9. All the secreted proteins with differential abundance between WT and RG1.9 are described in details in Table S1. **(C)** Functional classification of differentially abundant proteins based on their biological function as described in (54). The ratio of the abundant proteins corresponding to a specific functional category over the number of over-represented proteins in the specific strain is plotted. Dataset is filtered to secreted proteins, and cytoplasmic contaminating proteins are removed.

The identified secreted proteins of differential abundance compared to TK24 fell into six main functional classes (Fig. 3C and S4). Out of these classes, proteins of housekeeping functions were oversecreted in all the mutated strains RG1.9 (Fig. 3C), RG1.4 and RG1.5 (Fig. S4). In contrast, multiple hydrolases and metabolic process-related proteins were only secreted more in RG1.9, which showed a more severally affected phenotype (Fig. 3C and S4; Table S4).

As RG1.9 showed the highest number of proteins with differential abundance compared to TK24, we focused on this mutant for subsequent detailed experiments. Our LC-MS/MS analysis for the secretome of RG1.9 revealed that 13 proteins with cell wall-related functions and a protein with oxidation-reduction activity were oversecreted in RG1.9, for example: the Sec-pathway substrates: D-alanyl-D-alanine dipeptidase (D-Ala-D-Ala dipeptidase) (D6EQU5), a protein with lysozyme-like domain (IPR023346) (D6EKB6), a secreted protein with lysozyme-like (IPR023346), LysM (IPR018392) domains (D6ETM1) and Gamma-glutamyltransferase (D6EID2) with > 1.6 to 3- fold-change (Table S3 and S4) and the Tat-pathway substrates (beta-N-acetylhexosaminidase (A0A076M9V2) and L, D-transpeptidase (D6EYE3) with >1.8 and 2.9 fold change, respectively (Table S4).

Additionally, RG 1.9 secretome showed an oversecretion of 40 proteins with hydrolase activities. Collectively, these observations are suggestive of significant effects on the secretome caused by deleting secondary metabolite clusters on the *S. lividans* secretome.

### A reduced genome strain displays altered gene transcription

The differences seen in the protein abundance of the native secretomes of strains that carried secondary metabolite gene cluster deletions resulted from unknown alterations in metabolic regulation. We considered three possible mechanisms at play: a. elevated transcription of certain secreted protein genes, b. a change in levels of secretion apparatus genes or c. changes in chaperone abundance that might affect secretion. To examine this, we analyzed the transcriptomes of TK24 and of RG1.9 cells grown in NB medium by RNAseq.

349 genes out of a total of 7306 showed differential expression between RG1.9 and TK24 with a signal intensity ratio (M-value) >1 and adjusted *p*-value <0.05 (See materials and methods) (Fig. 4; Table S6). Out of these significantly differentially expressed genes, 196 were upregulated in RG1.9. These included 132 cytoplasmic, 5 nucleoid and ribosomal, 40 membrane and 14 secreted protein-encoding genes.

**Fig. 4:**
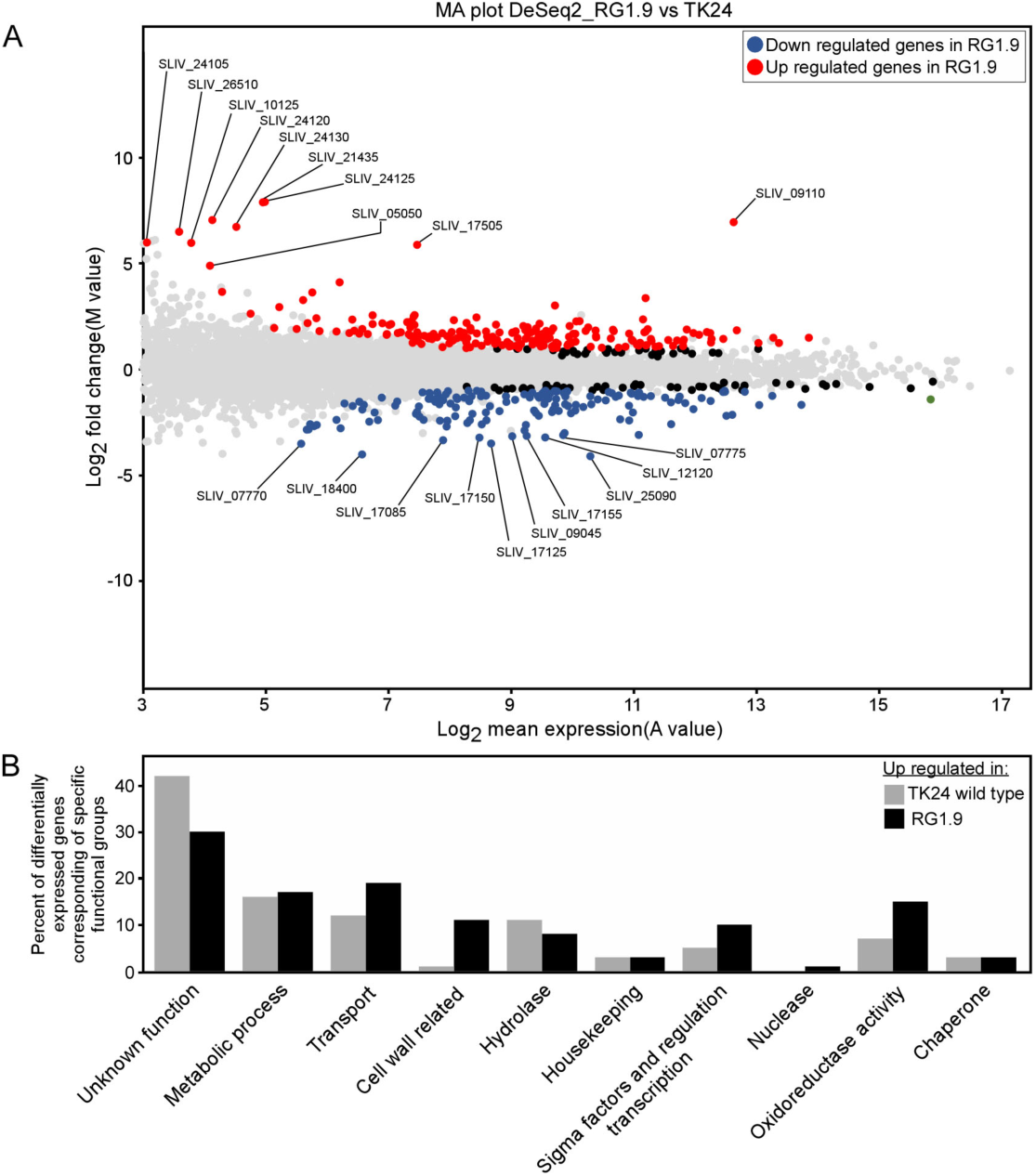
Differential expression of the reduced genome strain *S. lividans* TK24 RG1.9 against wild type TK24. **(A)** MA-plot for RNA-Seq datasets of TK24 wild type and RG1.9, which show up-regulated [M >1 and adjusted *p*-value<0.05, red circles, M(up)], down-regulated [M <-1 adjusted *p*-value<0.05, blue circles, M(down)] and non-regulated [M between +1 and -1 or/and adjusted *p*-value>0.05, grey circles, M(non)] genes. M-values (Y-axis) and A-values (X-axis) were calculated as indicated in the Material and Methods section. **(B)** Functional classification of differentially expressed genes based on their biological function as described in (54). The ratio of the differentially expressed genes corresponding to a specific functional category over the number of over-represented genes in the specific strain is plotted.

One obvious target for post-transcriptional regulation of secretion is the protein export machineries of the cell. For this, we monitored transcript levels of the export machinery components of the two main export systems in TK24, the Sec and the Tat, responsible for 93% and 7% of protein export, respectively (53). The gene encoding for the bifunctional preprotein translocase subunit SecDF, a component of the Sec export system, was upregulated 1.8-fold (Table S6) but none of the Tat-pathway component genes were affected.

Another group of proteins with a putative role in secretion are cytoplasmic chaperones that could bind selectively to groups of secretory proteins and regulate their secretion (55). Out of 58 genes encoding for cytoplasmic chaperones, 4 were significantly upregulated in RG1.9 (>1-fold increase and adjusted p-value<0.05). The gene that encodes for putative proteasome assembly chaperone 2 (IPR019151) (D6EKC1) showed the highest fold change (1.5-fold increase).

Similarly, the transcriptome of RG1.9 showed upregulation for 3 stress response-related chaperones: proteostasis-related stress response ATP-dependent serine protease Lon (D6EUK1)(56), thioredoxin reductase A (D6EMA7) that is related to the thioredoxin system (57) and nitrate reductase subunit delta (D6EHT8) with ∼1 to 1.4-fold increase in RG1.9 (Table S6).

Furthermore, differentially expressed genes in TK24 and RG1.9 were related to 10 functional groups (Fig. 4B; Table S6). 6 of these functional groups were upregulated in RG1.9, particularly those genes that related to oxidoreductase activity (18 genes) and cell wall-related function (20 genes) (Fig. 4B; Table S6). Additionally, 4 genes of the regulon of genes controlled by the cell envelope stress response Sig^E^ (58) were upregulated in RG1.9 (Table S6; showed with one asterisk). Similarly, 11 genes of the regulon of the redox hemostasis control Sig^R^ (59) were upregulated in RG1.9 (Table S6; showed with two asterisks).

Finally, we hypothesized that the presence of cell-wall stress should affect the transcription of genes with roles in nitrogen metabolism and nitrogen utilization for cell-wall biosynthesis, such as genes for the arginine biosynthesis pathway (60). RG1.9 transcripts showed upregulation for 4 genes of the arginine biosynthesis pathway *argH* (*SLIV_29875)*, *argC* (*SLIV_29825*), *argJ* (*SLIV_29830*) and *argG* (*SLIV_04065*), with 1.3-2.3-fold increase and adjusted *p*-value<0.05 (Table S6; showed with three asterisks).

These data suggest the presence of redox and cell wall stresses in TK24 resulted from secondary metabolite gene cluster deletion.

### A reduced genome strain displays metabolic alterations

To understand the difference in recombinant protein expression capacity between TK24 and RG1.9, we looked for differences in the exometabolomic profiles of both strains in NB medium, rich with amino acids which serve as carbon and nitrogen source for growth and protein formation. Free amino acid concentrations were analyzed for each strain in the mid-exponential, late-exponential and early stationary phases. To analyze the patterns in these time series data, a principle component analysis was performed. Two principle components explain 88% of the variance in the data (Fig. 5A). The metabolomic pattern diverges over time. Mid exponential data still cluster together for both strains, whereas later time points cluster together depending on the strain. Loadings of the amino acids (Fig. 5B) demonstrate how each variable (amino acid) influences the principal components. Aspartate, glutamate and asparagine strongly correlate and explain the clustering of the metabolomic footprint of both strains in the exponential growth phase. All three amino acids are rapidly consumed, and aspartate and glutamate are depleted after the mid-exponential phase (Table S7), as seen previously (61, 62).

**Fig. 5:**
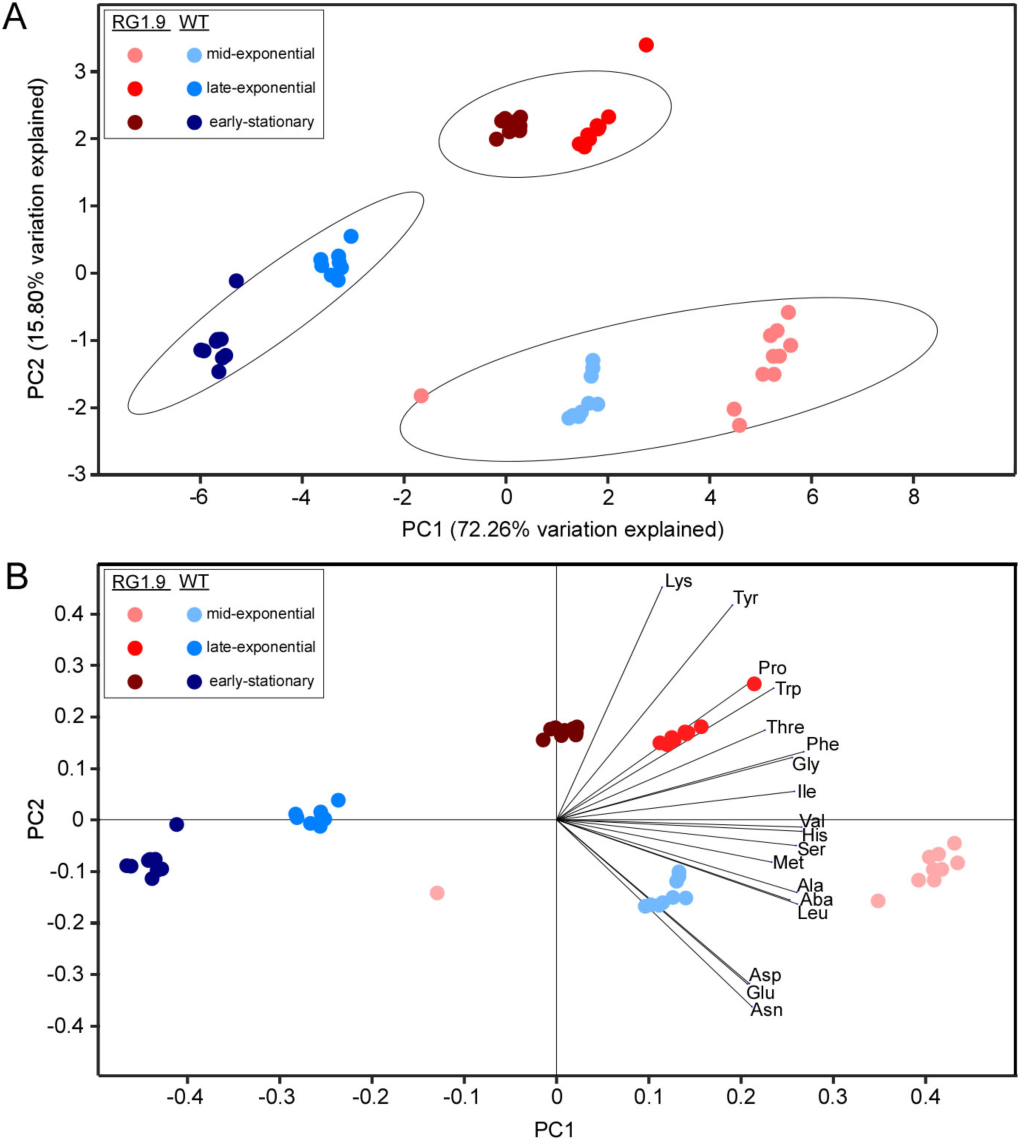
Exometabolomics changes between reduced genome strain *S. lividans* TK24 wild type and RG1.9. Principle component analysis was performed on amino acid concentrations in the medium measured in different growth phases**. (A)** PCA score plot with 95% confidence ellipses. (**B**) PCA biplot combining the score plot with loadings of the variables. Ala (alanine), Gly (glycine), Aba (α-amino-butyric acid), Val (valine), Leu (leucine), Ile (isoleucine), Thr (threonine), Ser (serine), Pro (proline), Asn (asparagine), Asp (aspartate), Met (methionine), Glu (glutamate), Phe (phenylalanine), Lys (lysing), His (histidine), Tyr (tyrosine), Trp (tryptophan) were used as variables and the experimental values as observations. Data from RG 1.9 and WT are printed in red and blue, respectively. Increasing color intensities corresponds to samples after 8 h (light ; mid-exponential phase), 16hrs (medium; late exponential phase) and 20hrs (dark; early stationary phase).

Lysine, proline, tyrosine and tryptophan are poorly correlated to the other amino acids, and concentration data show no uptake of these amino acids. Isoleucine, methionine, valine, histidine explain part of PC1 and threonine, phenylalanine and glycine can also be grouped and contribute to PC1 and PC2. Late exponential phase and early stationary phase seem to distinguish mostly for these two groups of amino acids. Although it is not possible to accurately estimate specific uptake rates, differences in uptake rates seem to exist between TK24 and RG1.9. The changed secondary metabolism in RG1.9 is most likely the explanation for the altered uptake patterns but it is not possible to retrieve exact correlations with metabolic pathways of secondary metabolism.

### Effect of deleted clusters on heterologous protein secretion

Finally, we tested the effect of genome reduction on the secretion of specific heterologous polypeptides. We first used a monomeric red fluorescence protein (mRFP) fused to a signal peptide as a model heterologous protein (3). The ORF of the mRFP gene was fused to the *vsi*-encoded signal sequence (*sp^SecV^-mRFP*) and cloned behind the *vsi* promoter in the high copy number plasmid pIJ486 to produce pIJ486/*sp^SecV^-mRFP* (3). The mRFP-expressing plasmid was transformed via protoplast transformation (see materials and methods) in TK24 and strains RG1.5 and 1.9, which secreted the highest amounts of total native secretome (Fig. 3A). To quantify the secreted mRFP, His-mRFP purified by metal affinity chromatography was used to generate calibration curves of fluorescence measurements against protein mass (3).

The amount of secreted mRFP varied between the mutated strains. mRFP secreted from RG1.5 (51 mg/g DCW), and RG1.9 (79 mg/g DCW) was more than what was secreted from TK24 (46 mg/g DCW) by 25 and 70%, respectively in NB at 24h (Fig. 6C; compare lane 3 and 5 with lane 1) while, at 48 h RG1.5 and RG1.9 secreted more mRFP than TK24 (by 8.5 and 24.5 %, respectively; Fig. 6C; compare lane 6 and 4 with lane 2). Analysis of the spent growth medium of TK24 and the derivative RG strains by SDS-PAGE, indicated a prominent band of ∼ 27kDa, close to the theoretical mass of mRFP (25.4 kDa) (Fig. 6A; filled arrow). Fluorescent measurement-derived amounts were close to those determined via semi-quantitative immuno-blotting using mRFP antibodies (Fig. 6B, compare lane 5 and 6 with lane 4).

**Fig. 6:**
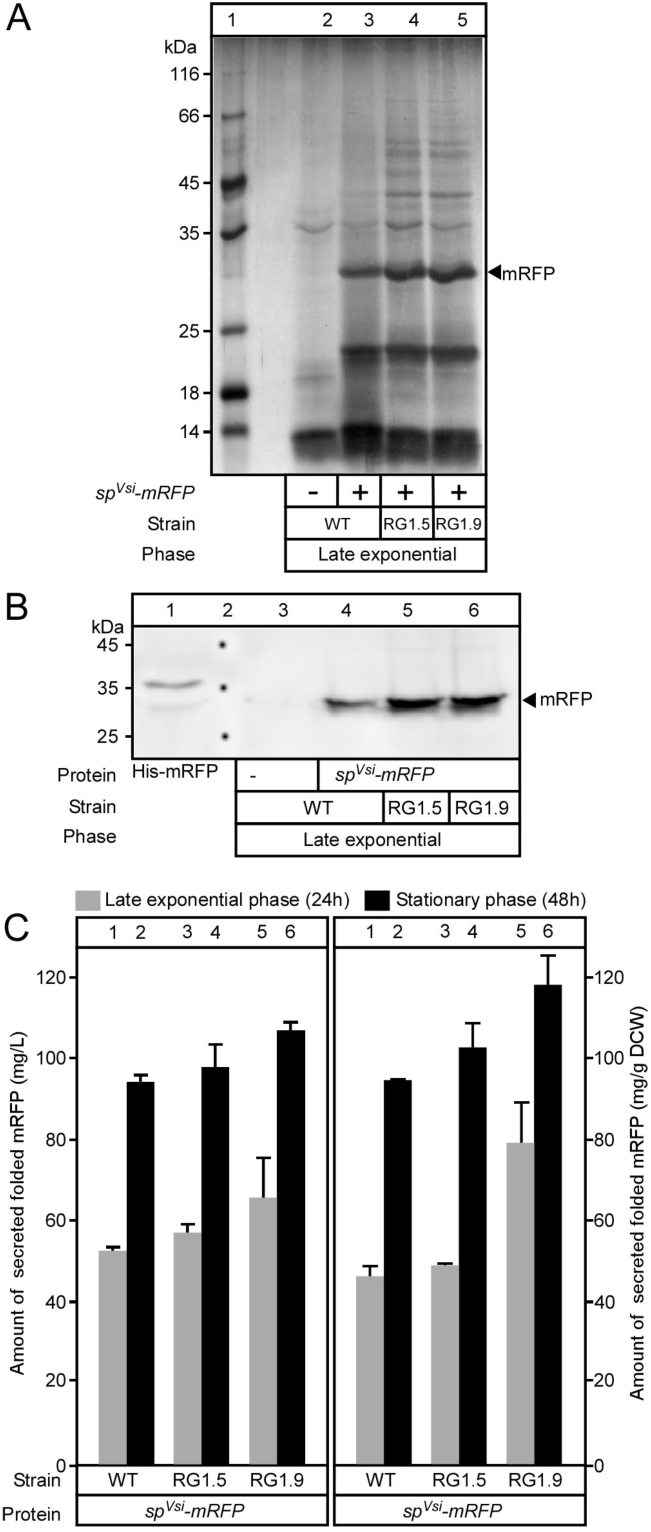
Effect of secondary metabolite genes clusters on heterologous proteins secretion. **(A)** Polypeptides from culture supernatants (2-16 µL/lane) that are equivalent to 0.1 mg correlated to a gram of DCW from TK24 or the reduced genome strains carrying a plasmid-borne copy of *sp^Secv^-mRFP* grown in nutrient broth (NB) for the indicated times were analyzed by SDS-PAGE and silver staining. Lane 1: Molecular weight marker as in (Fig. 3); Secreted mRFP (filled arrow) are indicated. **(B)** Western blots analysis for secreted mRFP nutrient broth (NB). Culture supernatants that are equivalent to 0.08 mg of mRFP of cells correlated to a gram of DCW of TK24 or the reduced genome strains carrying a plasmid-borne copy pf *sp^SecV^-mRFP* were separated by SDS-PAGE and visualized by western blotting using mRFP antibodies. **(C)** The amount of mRFP secreted (in mg/L and mg correlated to a gram of DCW) produced by TK24 or the reduced genome strains which were grown in nutrient broth (NB) for the indicated time related to its growth curves in the same media. *n*=3, values represent the mean ± SD.

Next, we tested the secretion of two other heterologous proteins in RG1.5 and RG1.9 compared to TK24. The heterologous gene encoding murine tumor necrosis factor (mTNFα) was cloned behind the *vsi*-encoded signal sequence and promoter as was mRFP to produce pIJ486/*sp^SecV^-mTNF* that was transformed in both the mutated strains and TK24 as previously described. To monitor and quantify the secreted mTNFα, calibration curves of purified His-mTNFα was generated using polyspecific antibodies α-mTNFα(5), respectively. Culture filtrates of TK24 and its derivative strains (RG1.5 and RG1.9) expressing *sp^SecV^-mTNF* in NB for 24h were collected and analyzed by SDS-PAGE followed by silver-staining (Fig. S5A) and α-mTNF immuno-staining (Fig. S5B). The quantification of mTNFα showed that it was secreted efficiently in RG1.5 and RG1.9 (195 and 230 mg/gDCW, respectively) measurably more than in TK24 (175 mg/gDCW) by ∼11% and 31%, respectively (Fig. S5C).

## Discussion

We aimed to explore whether genome reduction in *S. lividans* TK24 by knocking out non-essential secondary metabolite gene clusters affects the overall secretome and heterologous protein secretion. The results suggested the presence of complex system-level networks that control secondary metabolite production and protein synthesis and secretion.

Different combinations of secondary metabolite gene deletions affect the amounts of the TK24 secretome, while growth was only slightly affected (Fig. 2A). This would suggest that TK24 cells are under strict metabolic control. While the precise mechanism of metabolic regulation control remains unknown, this effect might be due in part to the multi-layered and cross-regulated networks of transcriptional regulators in TK24. As an example of how cross-regulated such networks are, the genes of the regulatory proteins for both the actinorhodin and undecylprodigiosine (ActII-ORF4 and RedD, respectively) are controlled by the same sigma factor *hrdD* (63). Additionally, *act*, *red* and *cda* clusters are transcriptionally downregulated by either the *wbl*A (SCO3579) (64) or a *tet*R family transcription regulatory gene (SCO1712) in *S. coelicolor* (65). Moreover, the promoter of *act*II-ORF4 is a target for at least 8 more different transcriptional regulator proteins: AdpA, LexA, DasR, DraR, AfsQ1, AtrA, ROK7B7 and AbsA2 (22). Disentangling these complicated cross-regulations will require extensive studies of RNAseq analysis combined with strains with carefully engineered deletions of such clusters of regulatory proteins and their common transcriptional activators and/or repressors.

The TK24 secretome changes quantitively and qualitatively by deletion of secondary metabolite gene clusters (Fig. 2, 3 and S3; Table S3 and S4). Besides these secondary metabolite clusters, strains that are missing either the sigma factor gene *hrdD* (RG1.5) or genes encoding for putative polysaccharide synthases *matAB* (*SLIV_22885* and *SLIV_22890*) (RG1.9) (Fig. 1; Table 1 and S1) secrete 2 to 3-fold more total polypeptides than does TK24 (Fig. 2A). These variations in polypeptide secretion amounts could be explained as results of strain responses to stresses caused by the deletion of these clusters in combination with either *hrdD* (58, 66, 67) or *matAB* (68, 69). In *S. coelicolor,* the closest relative of *S. lividans* TK24 (53), *hrdD* transcription is regulated by both sigma factor *sig^E^* (*SLIV_21000)* and *sig^R^* (*SLIV_12285*)*, which* are involved in sensing and responding to cell-wall (30), and thiol-oxidative (30) stresses, respectively. Furthermore, Sig^E^ controls the synthesis of both proteins that comprise the cell envelope (58) and secondary metabolite clusters, particularly actinorhodin (30). Consequently, *sig^E^* deletions resulted in actinorhodin overproduction in *S. coelicolor* (30, 70). Similarly, the deletion of the *matAB* genes in *S. coelicolor* resulted in forming a thinner wall lacking lamellae and patches, which caused cell wall stress (69). Collectively, cell wall stress might explain the elevated abundance of secreted proteins with cell wall-related function in the secretomes of a TK24 reduced genome strains, notably RG1.9 oversecreted >13 cell wall-related proteins (Fig. 5A and C; Table S4). Furthermore, RG1.9 transcriptome showed the upregulation of 24 genes of cell wall-related functions (four of them are part of the of Sig^E^ regulon (58)) (Table S6).

Additionally, the secretome of RG1.9 contained an over-abundance of multiple hydrolases, housekeeping proteins and proteins with metabolic process roles (Fig. 3C). These data may indicate the involvement of regulation other than the cell envelope stress in RG1.9 that affect polypeptide over-secretion. To have a better overview of such regulatory processes, we analyzed the transcriptome of RG1.9 and TK24 (Fig. 4; Table S6). The transcriptome of RG1.9 showed overexpression of many transcription regulator genes (e.g. sigma factors, transcription regulators and two-component systems (TCS)) (Table S6). Interestingly, out of these two-component systems, the genes encoding the TCS (a histidine-kinase sensor (*SLIV_09695*) and a response regulator protein (*SLIV_09690*)) were significantly upregulated in RG1.9 with >1.1 fold increase (Table S6). Propagation of this TCS enhances both antibiotic and protein secretion in *S. coelicolor* as they partially exert the function of the phosphorylated nucleotide stringent factor (ppGpp) (50, 71). Moreover, overexpression of this TCS homology in *B. subtilis* (*degS* and *degU*) elevates the secretion of extracellular degradative enzymes and controls the transition for the late exponential to the stationary phase of growth (71, 72). Regardless of transcription regulators, it is important to note that many other factors can affect protein secretion and are suggestive of more complex, hitherto unknown, mechanisms that act post-transcriptionally. These factors could involve the use of particular chaperones forming transient sequestering pools, as trigger factor is seen to do in *E. coli* (73), or as the quasi “SecB chaperone analogue” CspA may do in Gram-positive bacteria that have no SecB (55). Another important aspect of the secretion potential of the cell involves the secretion pathway components. In that context, the upregulation of *secDF* and 4 stress-related chaperones (Table S6) might be involved in RG1.9 elevated polypeptides secretion.

A functional classification for the differentially expressed genes in RG1.9 revealed the upregulation of 24 genes with cell wall-related function (4 of them are in the Sig^E^ regulon (58)) and 29 genes with oxidation-reduction related function (11 of them are in the Sig^R^ regulon (59)) (Fig. 4B; Table S6). These suggested the generation and involvement of redox stress in addition to cell envelope stress. The redox stress might be explained by either: (1) Actinorhodin cluster deletion. Actinorhodin contains quinone groups that can capture free electrons and act as antioxidants (74). Therefore, its deletion might cause oxidative stress; or (2) as cell stresses are known to occur concomitantly (75), some transcription regulators such as *hrdD* and *adpA* may be involved in coordinating these stress responses (76, 77).

RG1.9 showed a reduced biomass compared with TK24 (Fig. 2A). To understand the metabolic reasons behind this suppression, we determined the differences in the exometabolomic profiles of both strains. The metabolic footprint of both strains showed that aspartate, glutamate and asparagine were rapidly consumed, and depleted after the mid-exponential phase (Fig. 5B; Table S7) especially in RG1.9 that showed a fast depletion (Fig. 5B; Table S7). It is known that these three amino acids and glucose are among the favorite substrates for *Streptomyces* biomass production. Depleting these amino acids forces the cell to end the growth phase and enter the stationary phase (D’Huys et al., 2011; D’Huys et al., 2012). Furthermore, the rapid consumption of glutamate in RG1.9 could be correlated with cell-wall stress as it is a major structural component of *Streptomyces* cell wall (78). Notably, four genes of the arginine biosynthesis pathway (*argHCJ* and *G*), that forms arginine from glutamate and play a key role in cell-wall biosynthesis, were upregulated in the transcriptome of RG1.9.

The secretion of mRFP and mTNFα underline the suitability of *S. lividans* for heterologous protein production and secretion and corroborates previous observations with several other heterologous proteins (2, 3, 79, 80),(81). RG1.9 was the best secretor strain for mRFP and mTNFα, faring better than TK24 by 31 and 70%, respectively (Fig. 6 and S5). The data suggested a positive correlation between elevated native secretome and heterologous protein secretion. However, the changes in heterologous protein secretion efficiency were not limited to RG1.9 but are also seen in the wild type as well (2, 3, 79, 80),(6, 81). The molecular basis for these changes is unknown, but it might be due to the collective effect off the aforementioned factors.

In summary, deletion of secondary metabolite gene clusters and *matAB* cause redox and cell envelope stresses (Fig. 7). As a response to these stresses: (1) TK24 overexpresses, and oversecretes proteins with both cell-wall biosynthesis and remodeling and oxidoreduction activity functions as compensatory mechanisms (Fig. 7); (2) TK24 consumes elevated amounts of nutrients and components related to these functions (aspartate, glutamate and asparagine amino acids) (Fig. 7); (3) depletion of these amino acids forces TK24 to stop the growth and enters the stationary phase (Fig. 7). Growth suppression is negatively correlated with the over-secretion of polypeptides (2, 3, 30, 42, 53) (4). TK24 upregulates a two-component system (*SLIV_09695* and *SLIV_09690*) that enhances the secretion of extracellular degradative enzymes presumably to increase the availability of nutrients (Fig. 7); (5) Finally, TK24 overexpressed post-translatory regulators for the secretion process (SecDF and secretory chaperones) to regulate protein oversecretion (Fig. 7).

**Fig. 7:**
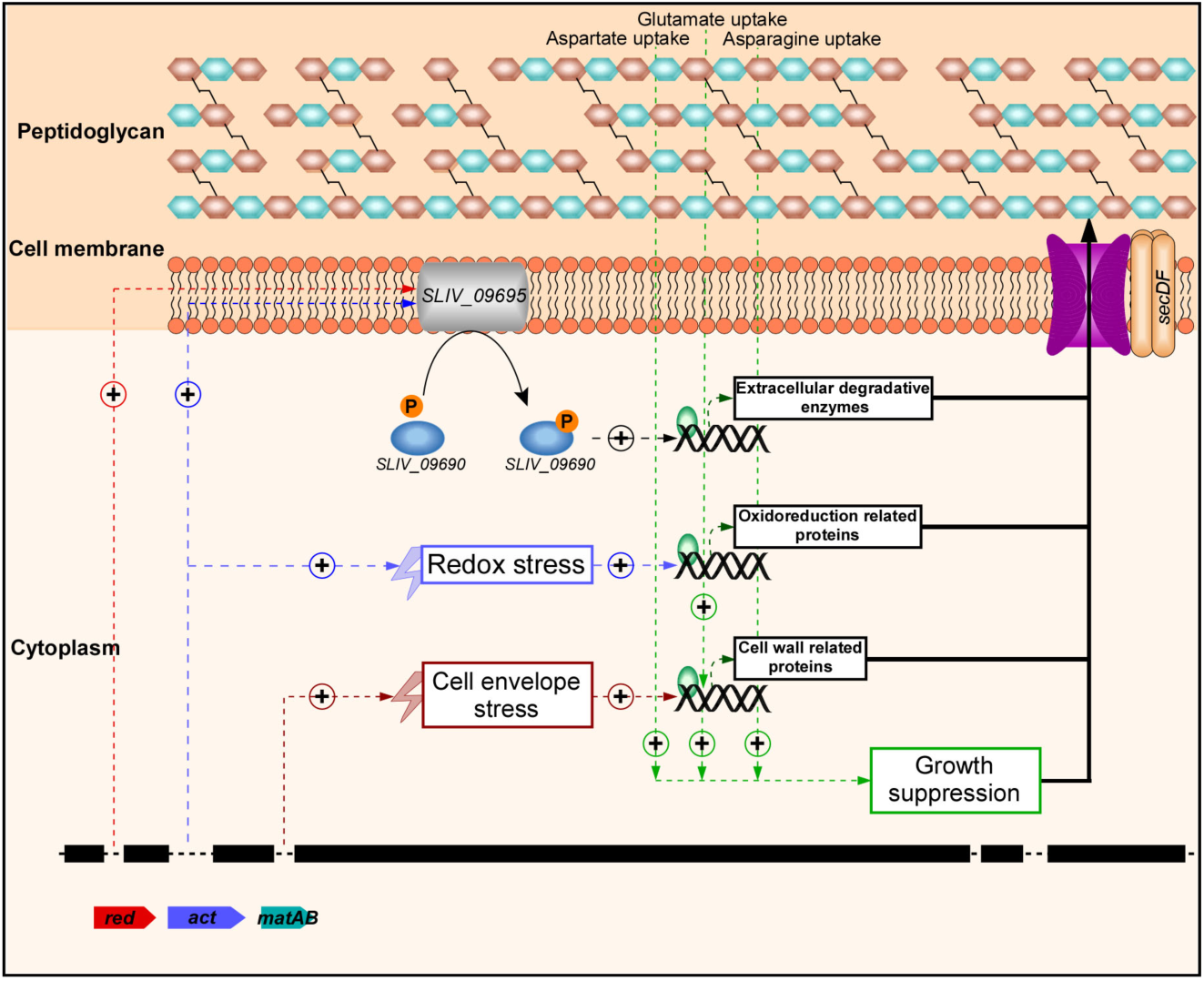
Proposed model for the effect of secondary metabolite gene clusters and *matAB* deletion on protein secretion. Deletion of both secondary metabolite gene clusters and *matAB* resulted in redox and cell envelope stresses. TK24 responds to these stresses by overexpressing and oversecreting protein with the cell wall and oxidoreduction-related function. Furthermore, TK24 rapidly consumes nutrients that are major components for cell wall remodelling (aspartate, glutamate and asparagine amino acids). Depletion of these amino acids forces TK24 to stop the growth and starts the stationary phase. Growth suppression is correlated with enhanced protein secretion. Additionally, TK24 upregulates a two-component system (*SLIV_09695* and *SLIV_09690*) that positively elevates extracellular degradative enzymes’ secretion for more nutrients availability. Moreover, TK24 overexpresses post-translational regulators (*secDF* and chaperones) to balance the oversecretion process. (+) indicates positive regulation.

This study provides an in-depth view of the complex regulation of secondary metabolite aspects and protein secretion in bacteria using a Gram-positive model cell. The combination of proteomics, metabolomics and transcriptomics (82, 83) with protein secretion (84) have revealed new processes and interactions that contribute towards deeper understanding of this fascinating complex process and its mechanistic regulation.

## Materials and Methods

### Strains, media and vectors used in the study

*Streptomyces lividans* TK24 was used as a wild-type strain (2, 85). *Streptomyces* cultures are grown at 27-30°C. Protoplast formation and subsequent protoplast transformation of *S. lividans* TK24 were carried out as described (86, 87). Deletion of secondary metabolite genes clusters from *S. lividans* TK24 to form reduced genome strains (RG) was carried out as described (41, 43).

Media used in this study were as described (2): Phage medium (88) (per liter: 10 g glucose, 5 g tryptone, 5 g yeast extract, 5 g Lab Lemco powder, 0.74 g CaCl_2_.2H_2_O, 0.5 g MgSO_4_.7H_2_O, pH: 7.2), Minimal Medium (MM) [per liter: 10 g glucose, 3 g (NH_4_)_2_SO_4_, 2.6 g K_2_HPO_4_, 1.8 g NaH_2_PO_4_, 0.6 g MgSO_4_.7H_2_O, 25 mL minor elements solution (per liter: 40 mg ZnSO_4_.7H_2_O, 40 mg FeSO_4_.7H_2_O, 40 mg CaCl_2_, 40 mg MnCl_2_.4H_2_O)], Minimal Medium with 5 g Bacto casamino acids/L (MM_C5_) and Nutrient Broth (NB) without NaCl [per liter: 8 g Nutrient Broth pH 6.9, i.e. per L: 5 g peptic digest of animal tissue, 3 g beef extract)]. For solid medium, MRYE (87) was used supplemented with the appropriate antibiotics, when necessary.

The productions of monomeric red fluorescence protein (mRFP), thermostable cellulase A from *Rhodothermus marinus* (CelA), a xyloglucanase from *Jonesia* sp. (Xeg) and active trimeric murine tumor necrosis factor alpha (mTNFα) in *S. lividans* TK24 and reduced genome strains were achieved As previously described in (2, 3, 5, 7)

### Growth conditions

*S. lividans* TK24 and its derivatives were precultured in 50 ml Phage medium (88) supplemented with thiostrepton (10 µg/ml) if necessary, and grown at 28 °C with continuous shaking (240 rpm; 48 h). After growth, the optical density of the preculture was measured at 600 nm (OD_600_) and the mycelia were harvested by centrifugation (3800 x g; 15 min; SIGMA 3-16K centrifuge) and washed twice with sterilized water. After homogenizing the mycelium in 50 mL of sterilized water, the strains were inoculated into 250-mL Erlenmeyer flasks containing 100 mL of nutrient broth medium (NB). In order to have the same amount of mycelia per each inoculum this equation was used: the volume of inoculum= (Final volume of culture X 0.25)/ OD_600_ (89). The flasks were shaken at 240 rpm and 28 °C, and pH was controlled using 100 mM MES buffer (pH 6.9).

### Dry cell mass determination

To quantify the dry cell weight (DCW), 10 mL of culture was centrifuged at 3800 x g for 15 min (SIGMA 3-16KL refrigerated centrifuge). The bacterial pellets were harvested, resuspended in sterilized water and filtered under vacuum using a 0.2 µm pore size filter (predried and preweighted; PORAFIL^®^ MV; Macherey-Nagel). The filter was once more dried (overnight 12-24 h in an oven at 60⁰C) and weighted for DCW determination.

### Fluorescence assays and quantification of fluorescent proteins

Fluorescence measurements were carried out in an Infinite® M200 microplate reader (Tecan) (mRFP: excitation at 550 nm/emission at 580 nm). To compare strain performance, mRFP fluorescence intensities, obtained at the transition to stationary phases were identified and evaluated. Error propagation was applied to calculate DCW-specific mRFP production.

The quantification of mRFP was carried out by using purified His-mRFP by metal affinity chromatography and generating calibration curves of both fluorescence measurements and protein mass (3). The amounts of mRFP detected via fluorescence assays were compared with the amounts quantified via western blotting mRFP antibodies (see western immunoblotting analysis below).

### SDS-PAGE and western blot analysis

Extracellular protein fractions of cultures of *S. lividans* and its derivatives were obtained by centrifugation (10 min, 4200 x g, 4°C). Precipitation of the proteins in the supernatant was, where applicable, carried out with trichloroacetic acid (TCA) (final concentration of 20% w/v; 4 °C). Proteins were separated by SDS-PAGE and as a standard, the Precision Plus Protein^TM^ Standard (All Blue) from Bio-Rad was used (90, 91). Proteins were visualized by silver staining or by Western blotting and immuno-detection with specific antibodies in combination with a suitable secondary alkaline phosphatase-conjugated antibody (Sigma). mRFP, CelA, and mTNFα antibodies were generated against lab-purified proteins at Davids Biotechnologie, Germany. Detection was carried out using the GE Healthcare Amersham ECL reagents and ImageQuant LAS 4000 Imager. High-resolution images were processed using ImageJ as described in (2).

### Secretomics sample preparation and measurement

Cells were removed by centrifugation (4,500 x *g*; 5 min; 4 °C) and subsequent filtration (syringe filter, 0.2 µm, cellulose acetate). Proteins contained in culture supernatants were precipitated via 25% v/v TCA precipitation (4 °C; 20 min). Precipitated proteins were pelleted via centrifugation (20,000 x *g*; 20 min; 4 °C), on a bench-top centrifuge. The pellet was washed twice with ice-cold acetone and re-pelleted via centrifugation (20,000 x *g*; 20 min; 4 °C). The protein pellet was then solubilized in 8M Urea in 1M ammonium bicarbonate solution (ABS). Polypeptide concentrations were measured using the Bradford reagent. Polypeptides (3 µg) were separated by 12% SDS-PAGE and visualized by silver staining (Shevchenko et al. 1996).

### Analysis of secretomes by nanoLC-MS/MS

A volume corresponding to the secreted polypeptides derived from 3 x 10^6^ cells (usually a volume equivalent to 20-40 µL of the initial cell culture) was used for in-solution tryptic digestion. The protein solution was initially diluted into urea (2 M final concentration in 50 mM Ammonium bicarbonate solution (ABS), followed by reduction of cysteines with 1 mM DTT (45 min; 56 °C), alkylation using 10 mM Iodoacetamide (IAA) (45 min; 22 °C; dark) and digestion using 0.015 µg Trypsin for 1.5 µg protein (Trypsin Gold, Promega, Fitchburg, Wisconsin; ratio trypsin/protein 1/100; overnight; 37 °C). Digested peptide solutions, were acidified with trifluoroacetic acid (TFA) to pH<2, desalted using STAGE tips (92, 93), and stored lyophilized at −20 °C, until the MS analysis.

Lyophilized peptide samples were re-suspended in an aqueous solution containing 0.1% v/v formic acid (FA) and 5% v/v Acetonitrile (ACN) and analyzed using nano-Reverse Phase LC coupled to a QExactive Hybrid Quadrupole - Orbitrap mass spectrometer (Thermo Scientific, Bremen, Germany) through a nanoelectrospray ion source (Thermo Scientific, Bremen, Germany). Peptides were initially separated using a Dionex UltiMate 3000 UHPLC system on an EasySpray C18 column (Thermo Scientific, OD 360 µm, ID 50 µm, 15 cm length, C18 resin, 2 µm bead size) at a flow rate of 300 nL min^-1^. The LC mobile phase consisted of two different buffer solutions, an aqueous solution containing 0.1% v/v FA (Buffer A) and an aqueous solution containing 0.08% v/v FA and 80% v/v ACN (Buffer B). A 60 min multi-step gradient was used from Buffer A to Buffer B as follows [0–3 min constant (96:4), 3–15 min (90:10); 15–35 min (65:35); 35–40 min (35:65); 40-41 min (5:95); 41-50 min (5:95); 50-51 min (95:5); 51-60 min (95:5)].

The separated peptides were analyzed in the Orbitrap QE operated in positive ion mode (nanospray voltage 1.5 kV, source temperature 250 °C). The instrument was operated in data-dependent acquisition (DDA) mode with a survey MS scan at a resolution of 70,000 FWHM for the mass range of m/z 400-1600 for precursor ions, followed by MS/MS scans of the top 10 most intense peaks with +2, +3 and +4 charged ions above a threshold ion count of 16,000 at 35,000 resolution. MS/MS was performed using normalized collision energy of 25% with an isolation window of 3.0 m/z, an apex trigger 5-15 sec and a dynamic exclusion of 10 s. Data were acquired with Xcalibur 2.2 software (Thermo Scientific).

Raw MS files were analyzed by the MaxQuant v1.5.3.3 proteomics software package (Cox, Mann 2008). MS/MS spectra were searched by the Andromeda search engine against the Uniprot *S. lividans* TK24 proteome (taxonomy: 457428, last modified May, 2016, 7320 protein entries; (23) and common contaminants (e.g. trypsin, keratins). Enzyme specificity was set to trypsin and a maximum of two missed cleavages were allowed. Dynamic (methionine oxidation and N-terminal acetylation) and fixed (S-carbamidomethylation of cysteinyl residues) modifications were selected. Precursor and MS/MS mass tolerance was set to 20 ppm for the first search (for the identification of the maximum number of peptides for mass and retention time calibration) and 4.5 ppm for the main search (for the refinement of the identifications). Protein and peptide false discovery rate (FDR) were set to 1%. FDR was calculated based on the number of spectra matched to peptides of a random proteome database (reversed sequence database) in relation to the number of spectra matching to the reference proteome. Peptide features were aligned between different runs and masses were matched (“match between runs” feature), with a match time window of 3 min and a mass alignment window of 20 min. Protein quantification was performed using the iBAQ algorithm (94) through MaxQuant software. Differentially abundant proteins were selected using the t-test and by comparing the fold difference of average protein intensities between the samples. P-values were further corrected for multiple hypothesis testing error using the Benjamini-Hochberg method (95).Thresholds for the analysis were set to adjusted p-value < 0.05 and fold difference > 2. Functional characterization of proteomics results was performed after filtering the dataset only to secreted proteins, excluding cytoplasmic contamination, using proteome annotation as described in the SToPSdb (54) (www.stopsdb.eu). The percentage of differentially abundant proteins that match a specific term over the total differentially abundant proteins for each condition was plotted. Keywords were derived after manual curation of the proteome.

### RNA isolation and genes differential expression

Samples for transcriptomics data analysis were taken during the late-exponential growth phase. The cells were grown in NB medium and the harvesting and RNA isolation was performed using Trizol reagent (Invitrogen) following the manufacturer’s instructions, and was treated with DNase I (Invitrogen) to remove chromosomal DNA contamination (96). Samples of 5 different biological replicates for each strain were isolated separately and pooled after quality control. The RNA quality was checked, and the TruSeq Stranded mRNA Library Prep Kit was done as described in (53, 97, 98).

Transcriptomics analysis and differential expression were carried out as described in (99). In brief raw FASTQ files were processed using the CLC Genomics Workbench (CLC Bio, Aarhus, Denmark). Raw reads were trimmed by their overall quality (score: 0.05; maximum ambiguous nucleotides: (2) and length (minimum length: 15 nucleotides). The filtered reads were mapped to *S. lividans* TK24 genome sequence, accession number (NZ_CP009124), with the default parameters (mismatch cost: 2; insertion cost: 2; deletion cost: 3; length fraction: 0.9; similarity fraction: 0.9; and ignore non-specific matches). The read counts were normalized using the DESeq-2 package in R (100).

Reads per kilobase per million mapped reads (RPKM) (101) were calculated based on the raw read counts per CDS. Differential gene expression analysis was performed using ReadXplorer v2.2 (102) using DESeq2 (100). The signal intensity value (A-value) was calculated by the average (log_2_) RPKM of each gene and the signal intensity ratio (M-value) by the difference of (log_2_) TPM. The differential RNA-Seq data was evaluated using an adjusted P-value cut-off of P ≤ 0.05 and a signal intensity ratio (M-value) cut-off of ≥ +1 or ≤ − 1 (fold-change of ±2).

### Metabolomics analysis

Exometabolomics analysis was performed on triplicate cultures of *S. lividans* TK24 wildtype and RS1.9 grown in nutrient broth. At several time points, free amino acids concentrations in the medium were measured. Cells were removed by centrifugation (4500 x g for 5 min) followed by microfiltration (syringe filter, 0.2 µm, cellulose acetate). Free amino acids were determined with the EZ:FAAST amino acid analysis kit of Phenomenex on a GC-FID (Perkin Elmer) according to supplier instructions. Measured amino acids are: alanine (ALA), glycine (GLY), α-amino-butyrate (ABA), valine (VAL), β-amino-isobutyric acid (BAIB), leucine (LEU), isoleucine (ILE), threonine (THRE), serine (SER), proline (PRO), asparagine (ASN), aspartate (ASP), methionine (MET), hydroxyproline (HYP), glutamate (GLU), phenylalanine (PHE), α-amino-adipic acid (AAA), glutamine (GLN), ornithine (ORN), cysteine (CYS), lysine (LYS), histidine (HIS), tyrosine (TYR), tryptophan (TRP). BAIB, HYP, AAA, GLN, ORN, and CYS were left out of the PCA analysis as these data were either too noisy or the amino acids were not really used as substrates. Principal component analysis was performed in MATLAB R2016b (The Mathworks) using included functions.

## Miscellaneous

Chemicals were from Sigma. Bacto Soytone from DIFCO Laboratories. DNA enzymes were from New England Biolabs and oligonucleotides from Eurogentec. Images were scanned in an ImageQuant™ 300 or a LAS4000 system and analyzed using the ImageQuant TL software to compare the intensity of the protein bands to those of the BSA calibration bands or other control proteins. SDS-PAGE and western blotting is described in the supplemental material. The mass spectrometry proteomics data have been deposited to the ProteomeXchange Consortium via the PRIDE partner repository (103) with the dataset identifiers under the same name of the manuscript using:

Password: dGvfqbBw

px-submission #642384

Submission Reference: 1-20230213-35563

## Authors’ contribution

HMB performed experiments of cell growth, protoplasts preparation and transformation, protein secretion and transcriptomics and proteomics data analysis and figures preparation; BT and KJ performed transcriptomics analysis; SK and BK design and performed metabolomics data; JK constructed reduced genome strains; VML supplied plasmids; AJ and CS analyzed data; EA and KS conceived, managed the project; HMB wrote the first draft of the paper; HMB and EA wrote the paper. All authors read and approved the MS.

## Acknowledgements

We thank Clara Marcelín-Leyva for preliminary experiments. This study was supported by: the European Union project (grant QLK3-CT-2002-02056, QLK3-LSHG-CT-2007-037586 and E.U.-FP7 project 613877 to JK, JA and AE); by the FWO (Fund for Scientific Research – Flanders); grant research project RiMembR (#G0C6814N) and by FWO/F.R.S.-FNRS ProFlow (“Excellence of Science - EOS” programme grant #30550343) to AE); RUN (#RUN/16/001 KU Leuven to AE); PROFOUND (W002421N; WoG/FWO) FOscil (ZKD4582 - C16/18/008; KU Leuven) to AE and SK): and by the Slovak Academy of Sciences (VEGA grant 2/00026/20 to JK). HMB was an Egyptian government doctoral scholar.

